# epitope1D: Accurate Taxonomy-Aware B-Cell Linear Epitope Prediction

**DOI:** 10.1101/2022.10.17.512613

**Authors:** Bruna Moreira da Silva, David B. Ascher, Douglas E. V. Pires

**Affiliations:** Systems and Computational Biology, Bio21 Institute, University of Melbourne,Melbourne, Victoria, Australia; Computational Biology and Clinical Informatics, Baker Heart and Diabetes Institute, Melbourne, Victoria, Australia; School of Computing and Information Systems, University of Melbourne, Melbourne, Victoria, Australia; The School of Chemistry and Molecular Biosciences, The University of Queensland, Brisbane, Queensland, Australia

## Abstract

The ability to identify B-cell epitopes is an essential step in vaccine design, immunodiagnostic tests, and antibody production. Several computational approaches have been proposed to identify, from an antigen protein, which residues are likely to be part of an epitope, but have limited performance on relatively homogeneous data sets and lack interpretability, limiting biological insights that could be derived. To address these limitations, we have developed epitope1D, an explainable machine learning method capable of accurately identifying linear B-cell epitopes, leveraging two new descriptors: a graph-based signature representation of protein sequences, based on our well established CSM (Cutoff Scanning Matrix) algorithm and Organism Ontology information. Our model achieved Area Under the ROC curve of up to 0.935 on cross-validation and blind tests, demonstrating robust performance and outperforming state-of-the-art tools. epitope1D has been made available as a user-friendly web server interface and API at http://biosig.lab.uq.edu.au/epitope1d.

## MAIN

B-Cell epitopes encompass a class of antigenic determinants that are dependent on the amino acids arrangement on the surface of the antigen protein structure. Contiguous stretches of residues along the primary sequence form linear epitopes, whereas non-adjacent residues, though nearby placed due to protein folding, form the discontinuous (or conformational) epitopes. Both forms impose a significant role upon the binding with its counterparts, the Immunoglobulins, that could be either in the form of membrane bound receptors or as antibodies, which are versatile macromolecules capable of recognising foreing threats [1], [2].

Identifying and selecting the appropriate epitope that could elicit an effective immune reaction in the host, and thus creating a protective memory immunity, is the fundamental basis of vaccine development [3], [4]. Being an extremely complex and multifactorial process, the amount of time spent in vaccine development is on average a decade and it can cost over 2 billion dollars for it to reach the market [5], [6]. Consequently, effectively aiding the selection of epitope candidates with computational techniques holds a promising role in this field in terms of substantial decreasing development time and cost.

Linear B-Cell epitopes account for only 10% among the two classes and although *in silico* prediction methods have significantly evolved over the past decades, varying from amino acid propensity scale scores [7]–[10], to combining physicochemical attributes and more robust machine learning techniques [11]–[15], their performance are still biassed towards specific data sets, leading to limited generalisation capabilities. A recently published approach [16], attempted to address these gaps by systematically cross-testing several previous benchmark data sets on their machine learning model and thus proposing two final models: a generalist and other specifically tailored for viral antigens. However, the general model was trained on data predominantly from HIV virus epitopes, which could be potentially non-representative, with the virus-specific model still presented modest performance, reaching a maximum MCC of 0.26 on blind-tests.

To fill these gaps, here we propose an explainable machine learning classifier based on the largest experimentally curated linear epitope data set so far, covering large span of organisms, presenting robust performance with different validation techniques, in addition to two new feature representation approaches: Graph-based signatures of protein sequences labelled with physicochemical properties, and Organism ontology identification of each input peptide, leveraging the classifier distinction between epitopes and non-epitopes.

## RESULTS

Two scenarios of evaluation were considered to assess the ability of epitope1D to accurately identify linear epitopes, based on the dataset employed and direct comparison with previous methods. The first comprises the use of well-established benchmarks: the BCPred set assessed, assessed under 10-fold cross-validation, followed by external validation with different independent blind test sets, as previously described (ABCPred-1, ABCPred-2, AAP, LBtope, iBCE-EL-1 and iBCE-EL-2). This scenario impartially compares the performance of our models with recent developments and identifies feature importance and their current limitations.

The second scenario consists of our newly curated, large-scale data set extracted from the IEDB database which includes organism information. Data sets filtered at different sequence similarity levels were employed: with internal validation evaluated using 10-fold cross-validation, and models assessed externally via blind tests. Furthermore, recently development methods, including BepiPred-3.0 [34], EpitopeVEC and EpiDope, also had their performance assessed on the same blind test to determine differences in performance.

### Comparison with alternative methods using the BCPred data set

#### Feature Representation: What makes up a linear epitope?

The data set extracted from BCPred, composed of 1,402 peptide sequences with a balanced class ratio of 1:1, was used to train and test several supervised learning models as a classification task. Their ability to distinguish between epitopes and non-epitopes was assessed and most predictive features identified. Outstanding features identified in this scenario include: (i) the maximum and minimum value of Antigenicity ratios in terms of amino acid triplets (AAT, measuring how overrepresented some amino acid are in the epitope class of this data set); (ii) the Composition pattern (CTD) of physicochemical and structural properties (hydrophobicity, normalised van der Waals volume, polarity, secondary structure and solvent accessibility); and (iii) the Graph-based signatures using both types of labelling: physicochemical properties (Acidic, Apolar, Polar Neutral, Basic and Aromatic) and Parker hydrophilicity prediction scale.

Using the interpretable classifier, EBM, to understand feature importance (**Figure S1** of Supplementary Materials), we observed that the antigenicity ratio features were in the top three most relevant: the maximum value within a peptide sequence (AAT_max), the interaction amid the maximum and the minimum (AAT_max x AAT_min) and the minimum value (AAT_min); followed by its interaction with specific Graph-based and Composition physicochemical descriptors, such as Apolar:Aromatic-8 (pairs of apolar and aromatic amino acids within a sequence distance cutoff of 8) and the amino acid composition in terms of Hydrophobicity (G1:Polar, G2: Neutral, G3: Hydrophobicity).

Further exploring interpretability, **Figure 3** depicts the rationale behind the model’s decision considering only the top most significant feature, the maximum AAT value, the cumulative sum of the antigenicity ratio scale for all possible amino acid triples within a peptide sequence. In the top chart, the horizontal axis details the feature range values, while the vertical axis shows the class, with the decision mark between the two classes set to 0 (above zero a higher probability of being an epitope and non-epitope otherwise). A clear decision point has been learned when *AAT_max* ranges from 4 to 6 (more precisely, larger than 5), which is also the average value of this feature for this data set. A possible interpretation of this result, in terms of the data set that includes 20-mer peptides only, can be that if at least 5 combinations of amino acid triplets are overrepresented in the sequence, there is a higher chance of it being an epitope. The bottom chart of the figure depicts the feature value distribution.

**Figure 3.**
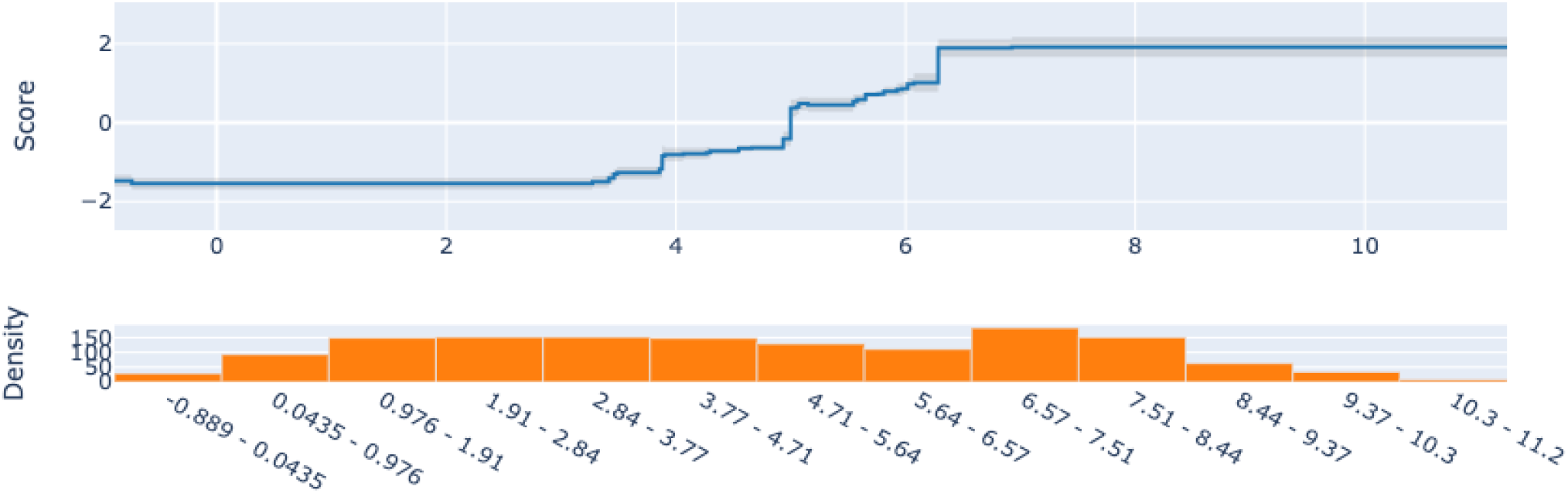
Behaviour of the feature AAT_max, which represents the maximum rate of Antigenicity for amino acid triplets, over its possible values ranging from -0.889 to 11.2. The epitope class probability increases when the AAT_max becomes larger than 5 (Score above 0). The bottom chart depicts the distribution of the corresponding data points in each feature interval.

#### Machine Learning Models

Under 10-fold cross-validation, the best performing models include Random Forest (RF) and Explainable Boosting Machine (EBM), both reaching an MCC of 0.72 and AUC of 0.92 and 0.93, respectively. Similar performance was observed using 5-fold cross validation. Table 1 provides a comparison of epitope1D with the previous methods that employed this data set including BCPred and EpitopeVec. Both methods used a Support Vector Machine (SVM) approach.

**Table 1.** Performance comparison of epitope1D using two different algorithms (EBM and RF) with BCPred and EpitopeVec methods under 10-fold cross-validation using the BCPred data set.

| METHOD | MCC | ROC-AUC | F1 |
| --- | --- | --- | --- |
| BCPred | 0.360 | 0.758 | —<sup>a</sup> |
| EpitopeVec | 0.620 | 0.880 | 0.810 |
| epitope1D (RF) | 0.720 | 0.920 | 0.850 |
| epitope1D (EBM) | **0.720** | **0.930** | **0.860** |

<sup>a</sup> Metric unavailable in original publication.

**Table 1.** Description of the data sets applied to train and evaluate epitope1D. The first column, named “Data Set”, is the name we are referring them to throughout the text; “Original Method” is where the set originally is derived from; “Epitopes’’ and “Non Epitopes’’ correspond to the total amount of labelled data within the set; “Experimentally Defined” indicates if the data from the two classes were experimentally assessed; “Data Source’’ specify the database from which the set was extracted; “Peptide Length” indicate the size of the peptides within the data set (specify the length -if fixed; or Assorted); “Validation” column designates if we apply the set for cross-validation or blind-testing purposes.

| Data Set | Original Method | #Epitopes | #Non Epitopes | Experimentally Defined | Data Source | Peptide Length | Validation |
| --- | --- | --- | --- | --- | --- | --- | --- |
| **General Benchmark Data Sets** |  |  |  |  |  |  |  |
| BCPred | BCPred | 701 | 701 | Only Epitopes | Bcipep and SwissProt | 20 mer | Cross-validation |
| ABCPred-1 | ABCPred | 700 | 700 | Only Epitopes | Bcipep and SwissProt | 20 mer | Blind |
| ABCPred-2 | ABCPred | 187 | 200 | Only Epitopes | Bcipep/SDAP and SwissProt | Assorted | Blind |
| AAP | AAP | 872 | 872 | Only Epitopes | Bcipep and SwissPro | 20 mer | Blind |
| LBtope | LBTope | 7,824 | 7,853 | Yes | IEDB | 20 mer | Blind |
| iBCE-EL-1 | iBCE-EL | 4,440 | 5,485 | Yes | IEDB | Assorted | Blind |
| iBCE-EL-2 | iBCE-EL | 1,110 | 1,408 | Yes | IEDB | Assorted | Blind |
| **New curated large-scale benchmark data set** |  |  |  |  |  |  |  |
| epitope1D Training | epitope1D | 20,638 | 103,281 | Yes | IEDB | Assorted | Cross-validation |
| epitope1D Testing | epitope1D | 5,264 | 25,716 | Yes | IEDB | Assorted | Blind |

In order to externally validate our model and assess its generalisation capabilities, different blind test sets were presented to the epitope1D (EBM) model as detailed in Table 2 and in Table S2 of Supplementary Materials, where we can also examine the performance achieved by previous methods such as BepiPred, BepiPred-2.0, EpiDope and EpitopeVec. Significant performance differences were observed for the methods when trained and tested using different data sources (*i*.*e*., Bcipep and IEDB databases). For instance, with the AAP data set (originally derived from Bcipep) in the first part of the Table 2, epitope1D and EpitopeVec (both trained using data from Bcipep database) achieved higher performances (MCC of 0.815 and 0.770, respectively) compared to the other methods that were trained using data derived from IEDB database (iBCE-EL, BepiPred and EpiDope). Alternatively, when applying the iBCE-EL testing data set (extracted from IEDB), our model and EpitopeVec achieved lower values of MCC, 0.092 and 0.095, compared to the model trained on data from this source (which reached a MCC of 0.454). However, in this scenario, some models trained using the same data source (e.g. BepiPred, BepiPred-2.0 and EpiDope), did not perform well either.

**Table 2.** Performance comparison with previous methods using three distinct blind test sets: AAP, ABCPred-2, and LBTope.

| METHOD | MCC | ROC-AUC | F1 | ACCURACY |
| --- | --- | --- | --- | --- |
| *Data set: AAP* |  |  |  |  |
| iBCE-EL | -0.036 | 0.528 | 0.350 | 0.494 |
| BepiPred | 0.217 | 0.665 | 0.600 | 0.604 |
| BepiPred-2.0 | -0.021 | 0.424 | 0.400 | 0.493 |
| EpiDope | 0.061 | 0.559 | 0.350 | 0.507 |
| EpitopeVec | 0.770 | **0.958** | 0.880 | 0.883 |
| **epitope1D** | **0.815** | 0.907 | **0.909** | **0.907** |
| *Data set: ABCPred-2* |  |  |  |  |
| ABCPred | _<sup>a</sup> | _<sup>a</sup> | _<sup>a</sup> | 0.664 |
| AAP | 0.292 | 0.689 | _<sup>a</sup> | 0.646 |
| iBCE-EL | -0.227 | 0.501 | 0.320 | 0.434 |
| BepiPred | 0.132 | 0.627 | 0.570 | 0.566 |
| BepiPred-2.0 | 0.181 | 0.62 | 0.480 | 0.555 |
| EpiDope | 0.091 | 0.541 | 0.360 | 0.508 |
| EpitopeVec | 0.445 | 0.778 | 0.720 | 0.718 |
| **epitope1D** | **0.543** | **0.841** | **0.848** | **0.781** |
| *Data set: LBTope* |  |  |  |  |
| iBCE-EL | **0.135** | **0.619** | 0.39 | 52.20 |
| BepiPred | 0.092 | 0.566 | **0.55** | **54.57** |
| BepiPred-2.0 | -0.001 | 0.476 | 0.42 | 49.95 |
| EpiDope | 0.036 | 0.559 | 0.35 | 50.34 |
| EpitopeVec | 0.065 | 0.548 | 0.51 | 52.98 |
| **epitope1D** | 0.067 | 0.527 | 0.333 | 52.80 |

<sup>a</sup> Metric unavailable in original publication.

Regarding the machine learning process, this behaviour raises concerns of potential biases in these data sets or lack of representativeness. Both cases lead to a lack of generalisation and can occur due to a variety of reasons, some of which we might conjecture: (i) all previous benchmark datasets listed here were adjusted to a highly balance ratio for epitope and non-epitope classes, which do not represent the biological truth; (ii) Bcipep database, as originally stated, are predominantly composed of viruses (HIV virus predominantly), which induces an underrepresentation of other organisms; (iii) Truncation/Extension approaches adopted by part of the methods to define a fixed peptide length, change the originally validated epitope sequence and may impose a learning bias towards an artificial set; (iv) The use of non-experimentally validated sequences to populate the non-epitope class, strategy adopted by some of the benchmarks, could lead the machine learning model to learn from imprecise or even erroneous data.

### Performance on a newly curated benchmark data set from the IEDB database

#### Feature Importance and Organism-aware Predictions

To address the potentially unrepresentative nature of the data, we curated an experimentally validated, large scale data set, integrated with high-level taxonomy organism information that incorporates the three main superkingdoms: Virus, accounting for 83% of the data and enclosed in 8 classes (Riboviria, Duplodnaviria, Monodnaviria, Varidnaviria, Ribozyviria, Anelloviridae, Naldaviricetes, Adnaviria), followed by 15% of Eukaryota with 5 classes (Metamonada, Discoba, Sar, Viridiplantae, Opisthokonta); and 2% of Bacteria with 7 classes (Terrabacteria group, Proteobacteria, PVC group, Spirochaetes, FCB group, Thermodesulfobacteria, Fusobacteria), totalising 20 binary categories. Organism taxonomy information was included in the set of features previously used (detailed in Table S3 of Supplementary), composed of four main categories Graph-based signatures, AAT Antigenicity ratio, Composition features and Organism taxonomy.

To better understand individual feature contributions to model outcomes, a post-hoc analysis using the SHAP [35] was employed using the Random Forest model. The importance order of each descriptor in this scenario can be understood as a ranked summary depicted in Figure S2 of Supplementary. The Antigenicity ratio group, with AAT maximum and minimum values, are very predictive features with higher values strongly correlating with the epitope class. The next most important feature is part of the Composition group, charge.G3, denoting a higher number of negatively charged amino acids in the epitope class. The fourth most important feature was organism taxonomy, particularly the Riboviria, potentially showing what the model learned as a consequence of the class imbalance data, where 87% of the Riboviria sequences belong to the non-epitopes, thus correlating the epitope class with organisms other than Riboviria. Graph-based signature features also play an important role to the model decision (*e*.*g*., Apolar:PolarNeutral-9 feature), denoting pairs of residues (polar and apolar) far apart from each other in sequence, though contributing to the epitope class (particularly for Riboviria sequence, Figure S3).

#### Machine Learning Models

In this second analysis, EBM and RF classifiers were assessed and performed equally, with Random Forest presenting a slightly faster training time, the reason why it has been chosen. 10-fold cross validation was performed using the epitope1D training set, followed by the blind-test evaluation with the independent blind test. Performance of state-of-the-art methods BepiPred-3.0, EpitopeVec and EpiDope on the same blind test were compared. Table 3 outlines the performance metrics for the cross-validation and blind tests, with epitope1D reaching a MCC of 0.613 and 0.608, respectively, in contrast with the best performing alternative method, EpitopeVEC, only achieving up to 0.139 MCC and BepiPred-3.0 achieving -0.007. Although the EpiDope method was trained using data from the same source, IEDB, we did not perform a homology removal check on the test set to guarantee direct comparison and avoid data contamination.

**Table 3.** Performance metrics on cross-validation (CV) and blind test, using epitope1D data set. State-of-the-art methods for linear b-cell epitope prediction were also appraised using the blind test: BepiPred-3.0, EpitopeVEC and EpiDope.

| Method | MCC | ROC-AUC | F1 |
| --- | --- | --- | --- |
| epitope1D (CV) | **0.613** | **0.935** | **0.658** |
| epitope1D (**blind test**) | **0.608** | **0.935** | **0.654** |
| EpitopeVec (**blind test**) | 0.139 | 0.618 | 0.306 |
| EpiDope (**blind test**) | 0.051 | 0.610 | 0.024 |
| BepiPred-3.0 (**blind test**) | -0.007 | 0.586 | 0.290 |

To better visualise model performance and Sensitivity/Specificity tradeoff for all the methods on the blind test, ROC curves were created (Figure 6), for the data set filtered at different similarity level cutoffs. The epitope1D curve, displayed in red, reached a significantly better ROC-AUC value of 0.935, compared to 0.618 from EpitopeVEC in yellow, 0.610 from EpiDope in blue and 0.586 from BepiPred-3.0 in green (for a 95% similarity cutoff -Figure 6A). Further analysis with different cutoffs of 90%, 80% and 70% (Figure 6B-D, respectively) applied in the independent testing set are likewise depicted, demonstrating that epitope1D consistently and significantly outperforms all alternative methods, with a small decrease in performance, further highlighting its robustness.

**Figure 6.**
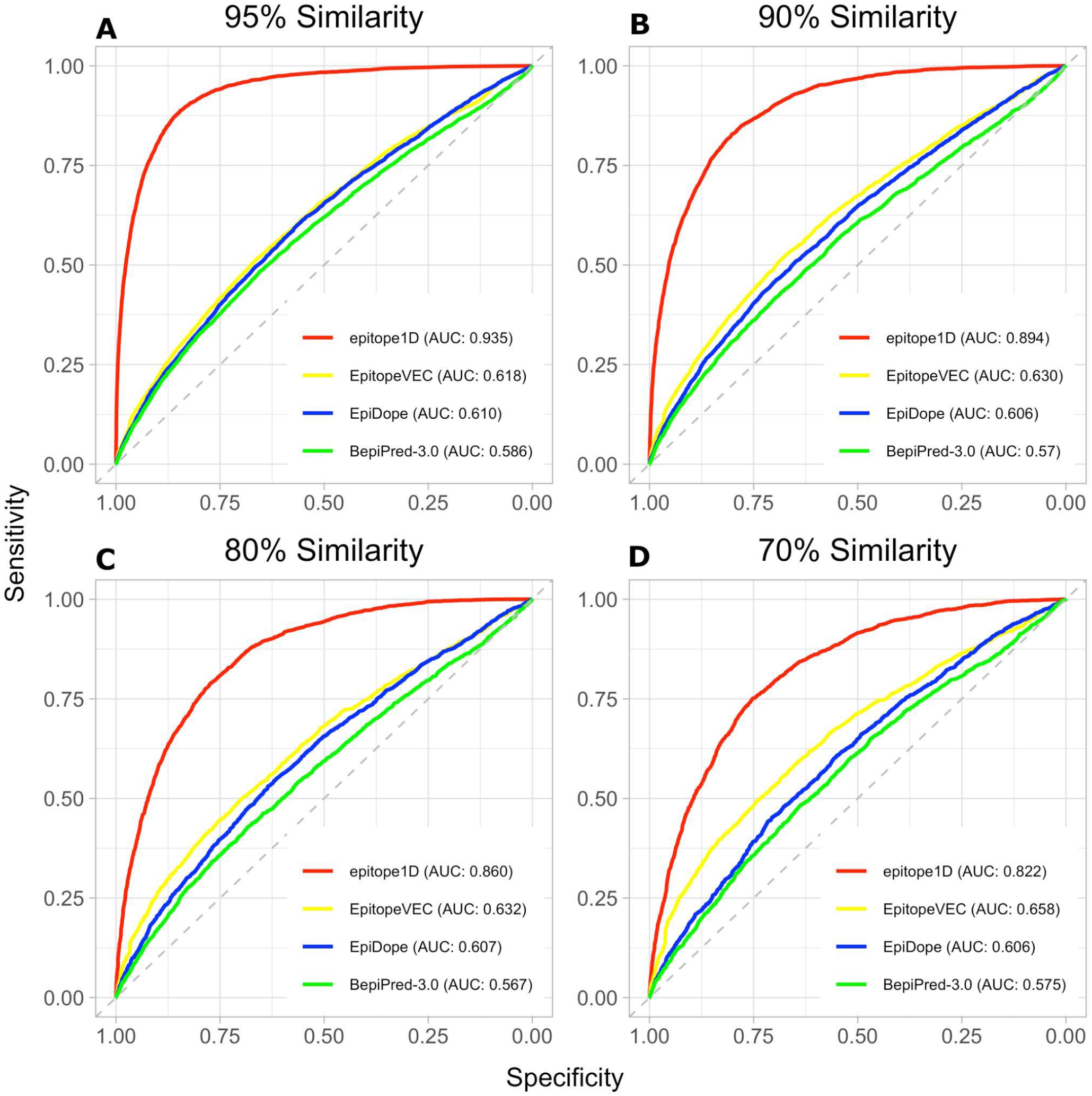
Performance comparison via ROC curves using the epitope1D test set at different similarity levels (95%, 90%, 80% and 70% for panels, A, B, C, and D, respectively). epitope1D achieved significantly higher AUC values of up to 0.935 (in red), followed by EpitopeVEC (in yellow), EpiDope (in blue) and BepiPred-3.0 (in green).

### Web server and API

epitope1D was made available as a friendly-user web server interface, where the user can input the protein or peptide sequence in fasta format, and select from a drop-down menu the equivalent organism taxonomy representation (Figure S4). In addition, an Application Programming Interface (API) enables for batch submissions and integration to standard analytical pipelines, contributing to reproducibility and usability of the resource.

## CONCLUSIONS

Linear B-cell epitope prediction is yet an extremely challenging task in which the sophisticated biological mechanism underlying the binding among Antibody-Antigen poses challenges to computational and experimental methods. The majority of previous benchmark data sets dated from up to 15 years ago, also suggesting a potential bias and lack of organism representativeness, leading to weak to poor generalisation capabilities.

epitope1D fills these gaps via an explainable machine learning method, built on the largest non-redundant experimentally validated data set to date, composed of over 150,000 data points, consisting of a diverse set of organisms within Virus, Eukaryota and Bacteria superkingdoms. epitope1D leverages well-established as well as novel features engineered to model epitopes, including a new graph-based signature to train and test taxonomy-aware and accurate predictors.

A comprehensive comparison of our method with state-of-the-art tools showed robust performance across distinct blind-test sets, with epitope1D significantly outperforming all methods, thus highlighting its generalisation capabilities. We believe epitope1D will be an invaluable tool assisting vaccine and immunotherapy development and have made it freely available to the community as an easy-to-use web interface and API at http://biosig.lab.uq.edu.au/epitopde1d.

## METHODS

epitope1D has four general steps, as can be seen in **Figure 1**: (1) Data collection and curation; (2) Feature Engineering: evaluation of current used features and new proposed features to represent peptide sequences; (3) Explainable Machine Learning: assessment of several supervised learning algorithms and further comparison among previous work using different data sources and explainability resources; and (4) Web server interface: development of an easy-to-use platform for end users.

**Figure 1.**
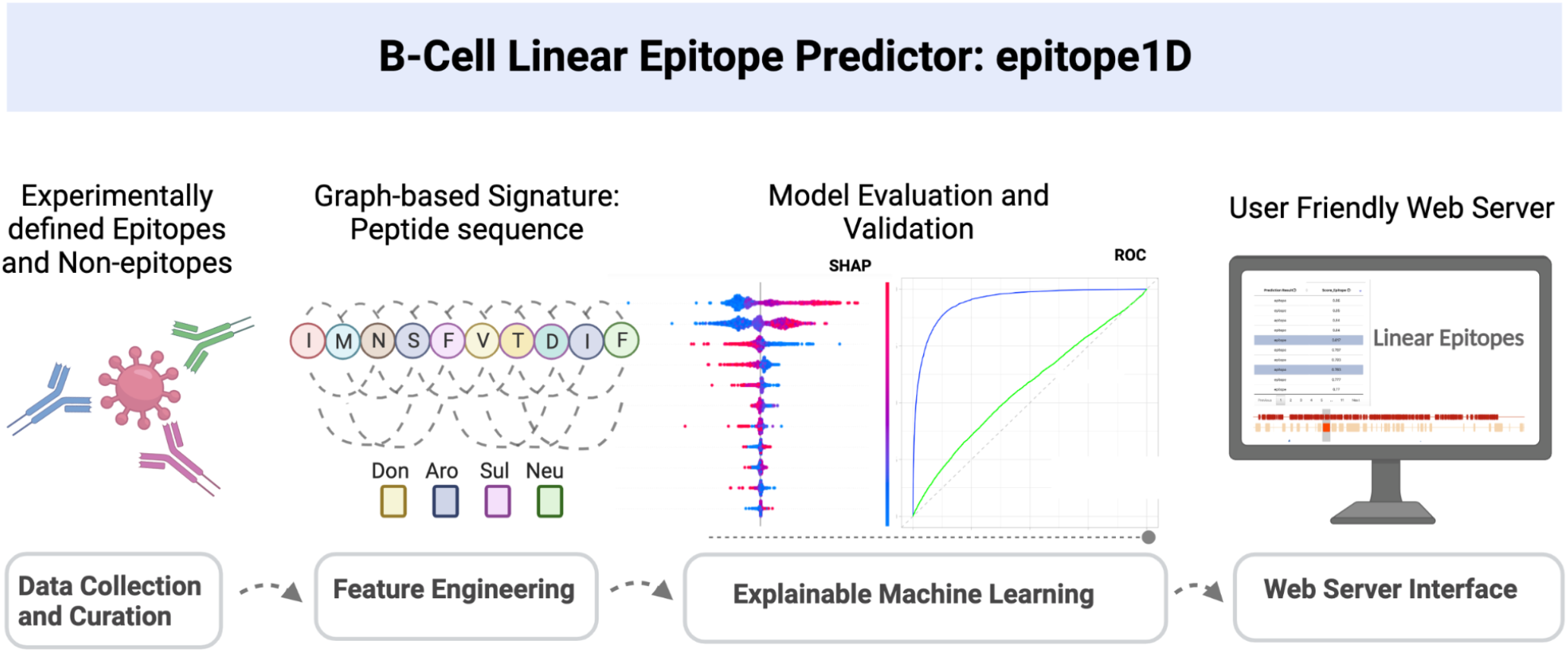
epitope1D workflow with 4 main stages: (1) Data Collection and Curation, which includes the selection of benchmarks and also the curation of an updated large-scale data set; (2) Feature Engineering, representing the step where all descriptors were calculated; (3) Explainable Machine Learning, in which the supervised machine learning classifiers were analysed in terms of their predictive power, explainability, and assessed via cross-validation and blind-test approaches; (4) Web Server Interface, where epitope1D is made publicly available as a user-friendly web interface and API.

### Data Collection

Well-established reference data sets were used to train, test and validate supervised learning algorithms, as a classification task. A comprehensive list of benchmark data sets, derived from ABCPred method [11], BCPred [17], AAP [18], LBtope [13], and iBCE-EL [12] was further employed to impartially evaluate the predictive power of our selected model against the original ones and state-of-the-art tools such as EpitopeVec [16], BepiPred [19] and BepiPred-2.0 [14]. A detailed review of them is located in Supplementary. Subsequently, given that most data sets were outdated (dated from 5 to 16 years ago), we have curated a newly updated data set derived from IEDB database [20], which will be used as the final basis for our model. Table 1 summarises the information of all data used in this work and further analysis can be read below.

### Curating a New Experimental Benchmark Data Set

We have curated a new set derived from the IEDB database aiming to reflect data availability on experimentally confirmed linear b-cell epitopes and also non-epitopes. Our main motivation arises because most of the previously mentioned data sets were collected more than 10 years ago and their negative class sets (non-epitope sequences) weren’t empirically proven. In addition, epitope sequences derived from the Bcipep database [21] can impose an obstacle to model generalisation, given that around 80% of them refers only to HIV virus. Therefore, organism information from each sequence in the new data is taken into account to assess whether the taxonomy can aid distinguish epitopes from non-epitopes, in a variety of subspecies.

The curation process of the new experimental data set comprises the following steps: (1) Download all possible (any host and disease) linear peptides of B-cell epitopes and non-epitopes as of June/2022; (2) Keep only the epitopes and non-epitopes confirmed in two or more different assays; (3) Consider solely the peptides with length between 6 and 25 amino acids, since 99% of linear epitopes range within these lengths [1], [12], [14]; (3) Remove sequences that were present in both classes simultaneously; (4) Exclusively retain entries that contain information about the source organism; (5) Perform a systematic sequence redundancy removal step using CD-HIT at different thresholds (95%, 90%, 80% and 70%) to assess the overall learning efficiency within high to medium similarity. Considering that a single antigen may contain several different epitope stretches that lead to distinct antibody bindings, it is worth accommodating the majority of available epitope sequences belonging to each antigen protein [2].

The final set, with a maximum of 95% similarity, is composed of 154,899 data points, in which 25,902 are epitopes, encompassing 1192 sub-species that were aggregated into a higher taxonomy parent organism lineage of 20 classes, each belonging to the superkingdom of Virus, Eukaryota or Bacteria. The final set was randomly divided into a training set with 123,919 data points, in which 20,638 are epitopes (ratio 1:6) and corresponds to 80% of the data, and the remain 20% as an independent test set with 30,980 data points with the same epitope/non-epitope proportion.

### Feature Engineering

To better characterise peptide sequences that might compose an epitope, previously used descriptors as well as novel features were evaluated. To reduce model complexity, a forward stepwise greedy feature selection algorithm was applied [22] to retain only the most representative set. The description of new proposed features is introduced here, while auxiliary features are described in Supplementary Materials. An overall summary is also provided in Table S1.

#### Graph-based signatures

We have designed a new graph-based feature, tailored for modelling linear epitopes of flexible length, inspired by the CSM algorithm [22]–[25]. The key idea was to model distance patterns among residues (nodes) at different distance cutoffs (each distance inducing the edges of a graph), which are summarised in two approaches: cumulative and non-cumulative distributions. Sequence graphs were labelled in two ways: (1) the corresponding scales of hydrophilicity prediction [8], beta turn prediction [26], surface accessibility [9] and antigenicity [10]; and (2) the amino acid physicochemical properties, such as Apolar, Aromatic, Polar Neutral, Acid or Basic, as done previously [22], [27], [28]. Figure 2 shows the steps comprising the new graph-based feature.

**Figure 2.**
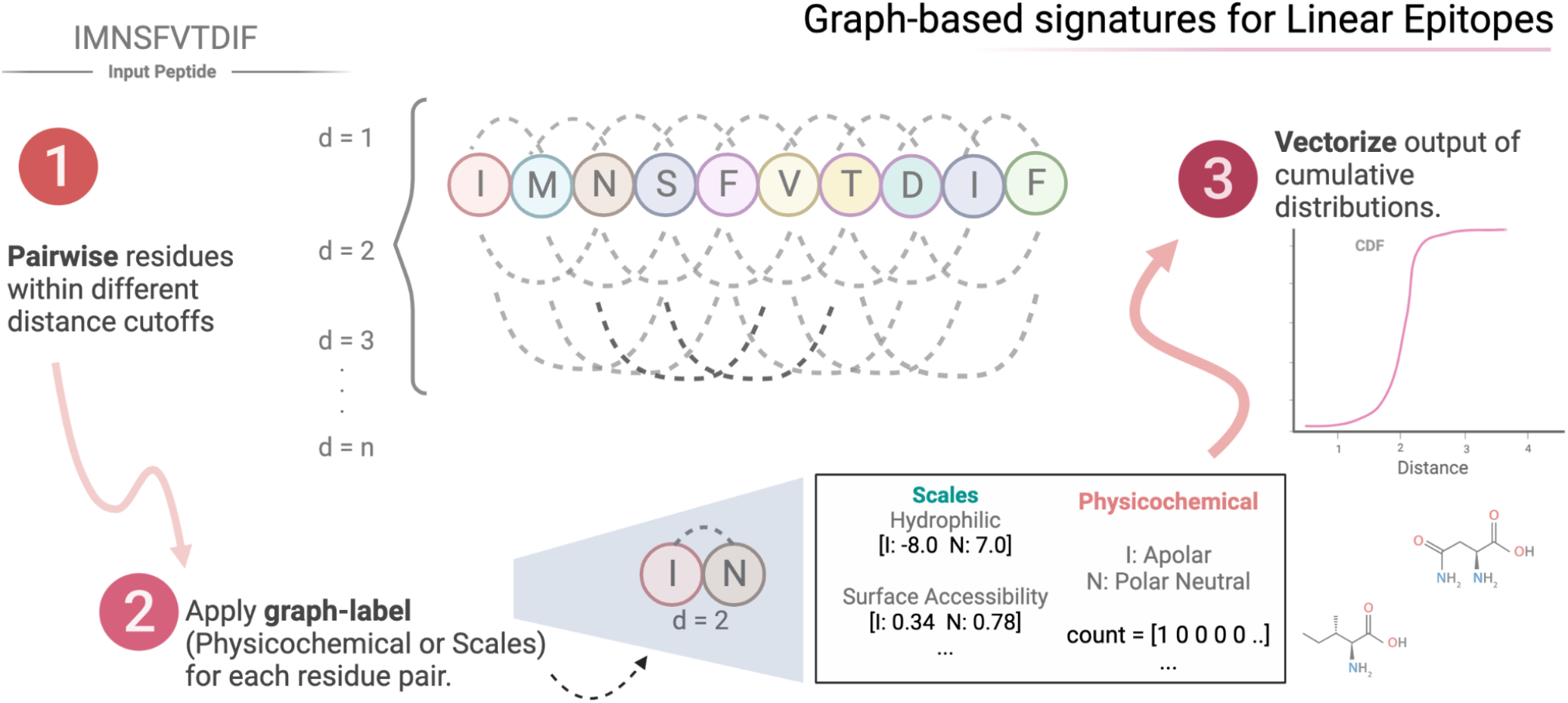
Modelling linear epitopes using graph-based signatures. The first step comprises the selection of residue pairs at incremental sequence distances and apply different types of labelling approaches on them. The third and final step is to vectorize the output as cumulative distances between different label pairs.

#### Organism Identification

The organism source information, extracted from the IEDB database together with each peptide sequence, is expressed by its ontology identifier deriving from two sources: (1) The Ontobee data server [29] for the NCBI organismal taxonomy; and (2) The Ontology of Immune Epitopes (ONTIE), which is an internal web resource from IEDB that was then converted back to the corresponding NCBI taxonomy term for standardisation. This information was used aiming to contribute with epitope identification addressing the pain point in the machine learning process that arises from high heterogeneity in organism classes [16], [30]. To transform the 20 ontological terms, described in Table S1, from categorical data into numerical, a one-hot encoding process was imposed. This descriptor was applied in the new curated benchmark data set only.

### Machine Learning Methods

As epitope identification could be described as a binary classification task, various supervised learning algorithms were assessed using the SciKit Learn Python library [31]. These included Support Vector Machine (SVM), Adaptive Boosting (Adaboost), Gradient Boosting (GB), Random Forest (RF), Extreme Gradient Boosting (XGBoost), Extra Trees, K-nearest neighbour (KNN), Gaussian Processes and Multi-Layer Perceptron (MLPC). In addition, an inherently interpretable method named Explainable Boosting Machine (EBM), a type of Generalised Additive 2 Model (GA2M) and considered as a glassbox model, was assessed via the open-source Python module InterpretML [32]. The goal of interpretable machine learning models is to provide a rationale behind prediction that would allow for meaningful biological insights to be derived, also assisting in the possible biases and errors as well as highly predictive features.

Performance evaluation for each model was done based on Matthew’s Correlation Coefficient (MCC), which is a robust statistical measure appropriate for imbalance data sets [33]. Complementary performance metrics were also used including F1-score, Balanced Accuracy and Area Under the ROC Curve (AUC). Performance between internal validation (k-fold cross validation) and external validation (blind tests) was contrasted to infer generalisation capabilities.

## SUPPLEMENTARY DATA

Supplementary data are available online at https://academic.oup.com/bib.

## DATA AVAILABILITY

epitope1D and associated data sets are available through a user-friendly and freely available web interface and API at http://biosig.lab.uq.edu.au/epitope1d, enabling seamless integration with bioinformatics pipelines and supporting quick assessment of sequences to support diagnosis and vaccine design.

