## Supplementary Materials for "epitope1D: Accurate Taxonomy-Aware B-Cell Linear Epitope Prediction"

#### General Benchmark  Data Sets

In total, we have selected seven benchmark data sets for comparison purposes, originally curated by previous methods.

- ABCPred curated two data sets, one for training and another for testing purposes. The first, that we named ABCPred-1, is composed of 700 experimentally defined epitopes retrieved from the Bcipep database [1] with a maximum length of 20 amino acids, and 700 peptide sequences of 20 amino acids long selected at random from SwissProt database [2] to represent non-epitopes. The epitope set may have been extended based on the original antigen sequence to achieve 20-mer size, as 90% of the epitopes from Bcipep have length smaller or equal to 20 amino acids. The second set, ABCPred-2, is composed of 187 epitopes sequences of different lengths, 59 from Bcipep, with no overlap with the first set, and others 128 were extracted from the Structural Database of Allergenic Proteins (SDAP) [3]. The non-epitopes are composed of 200 non-allergenic protein sequences of well-known allergen foods, from [4], that were a result of mining the SwissProt database. No systematic redundancy checks were reported for these data, only the removal of identical sequences.
- BCPred provides a non-redundant data set intended for cross-validation, with sequence similarity cutoff of 80% applying CD-HIT [5] analysis. It is composed of 701 experimentally defined epitopes which, besides having a sequence extension if lower than 20 amino acids, also were subjected to shortening, if longer, trimming both peptide ends to achieve a 20-mer. The negative class is composed of 701 20-mers randomly chosen from the SwissProt, certifying that none are part of the positive class.
- AAP method curated a set for internal cross-validation composed of 872 epitopes from the Bcipep database, in an analogous process to ABCPred in terms of sequence prolongation or reduction to achieve peptides with fixed 20-mer size, likewise keeping exclusively unique sequences. The negative class was again created by indiscriminately picking 20-mer sequence

fragments from the SwissProt database, guaranteeing that there was no overlap with the positive class.

- LBtope was the first method to curate a larger set from an experimentally assessed source, IEDB, for both epitopes and non-epitopes. A non-redundant set was built for internal cross-validation, based on CD-HIT with 80% similarity, composed of 7,824 epitopes and 7,853 non-epitopes with equal 20-mer size, applying extension and truncation on both classes based on the original antigen sequence.
- iBCE-EL also extracted experimental data from IEDB database, gathering a reduced set (70% homology threshold) of 5,550 epitopes and 6,893 non-epitopes, divided in two separated groups: 80% used for training purposes (4,440 epitopes and 5,485 non-epitopes) and 20% (1,110 epitopes and 1,408 non-epitopes) designated for independent evaluation.

#### Feature Engineering

**Amino acid composition and antigenicity scales.** The first group of features used for peptide representation was the amino acid composition, calculated using the iFeatures Python package [6] with different arrangements: (i) k-spaced amino acid pairs, which calculates the frequency of amino acid pairs apart from each other by a number of residues; (ii) Dipeptides; (iii) Tripeptides; and (iv) Enhanced, that measures the number of each possible amino acid over a sliding window of defined length within the sequence; (v) Composition, Transition, and Distribution patterns of physicochemical and structural properties - named CTD - calculated splitting the amino acid sequence in three groups according to the attributes: Hydrophobicity (G1:Polar, G2: Neutral, G3: Hydrophobicity), Normalised van der Waals volume (G1: volume range 0 - 2.78, G2: 2.95 - 94.0, G3: 4.03 - 8.08), Polarity (G1: polarizability value 0 - 1.08, G2: 0.128 - 120.186, G3: 0.219 - 0.409), Secondary structure (G1: Helix, G2: Strand, G3: Coil), Charge (G1: Positive, G2: Neutral, G3: Negative) and Solvent accessibility (G1: Buried, G2: Exposed, G3: Intermediate). In addition, AAindex [7] was also evaluated. The antigenicity scale in the form of amino acid pairs (AAP) [8] and in triplets (AAT) [9] were also assessed and represent the normalised ratio frequency of amino acid pairs or trios between epitope and non-epitope classes. Both scales were updated and recently evaluated by EpitopeVec, where the positive and negative class information were derived from the entire Bcipep database and SwissProt, respectively. Here, besides the average score values of AAT/AAP, we also assessed the cumulative sum (the maximum value) for each peptide sequence and the minimum value. Moreover, we have also extracted an additional AAT scale, but using IEDB database as source for positive and negative class, filtering the linear b-cell epitopes confirmed in two or more experimental assays with length between 5 and 30 amino acids.

#### BCPred data set: Interpretable analysis of EBM model on training

Taking the interpretable classifier, EBM, to understand the ranking feature importance, we can observe in **Figure S1** that the antigenicity ratio accounted for the top three in different combinations: the maximum value within a peptide sequence (AAT_max), the interaction amid the maximum and the minimum (AAT_max x AAT_min) and the minimum value (AAT_min); followed by its interaction with a specific Graph-based and CTD physicochemical descriptors, such as Apolar:Aromatic-8 (pairs of apolar and aromatic amino acids within a distance cutoff of 8) and the amino acid composition in terms of Hydrophobicity. The last four ranked features symbolise: the graph-based labelled with Parker hydrophilicity scale, Parker-9_min, which takes the minimum scale values within a distance of 9 residues, then an interaction between AAT maximum and the Graph-based average scale of Parker in distance 9, shown as AAT_max x Graph-Based (Parker-9_avg), and ends with the two Graph-based descriptors using pairs of aromatic and basic amino acids, within a distance cutoff of 7, and pairs of acidic and apolar within distance 1.

#### Performance on a newly curated benchmark data set from the IEDB database

To deepen our understanding about the Graph-based representation of amino acid pairs labelled as Apolar:PolarNeutral-9, the fifth most important descriptor ranked in the SHAP plot, we further identified that it correlates with the Riboviria Organism feature, as depicted in Figure S3. From this, we were able to infer that the more frequent these physicochemical pairs are, the higher the likelihood of the sequence being labelled as epitope.

### FIGURES

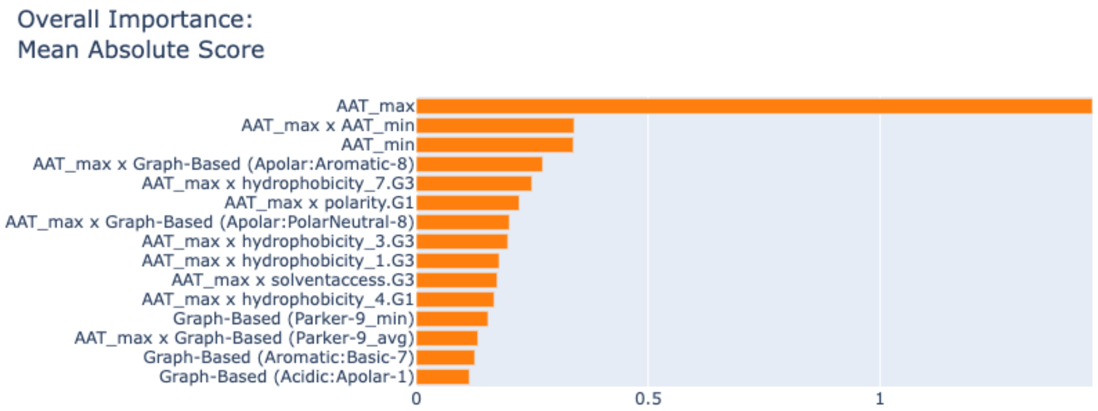


**Figure S1.** Feature importance ranking of EBM method using BCPred data set as training showing the Antigenicity ratio in multiple combinations (AAT maximum and minimum values), some Graph-based signatures and Composition patterns of Hydrophobicity (G1: Polar and G3: Hydrophobicity).

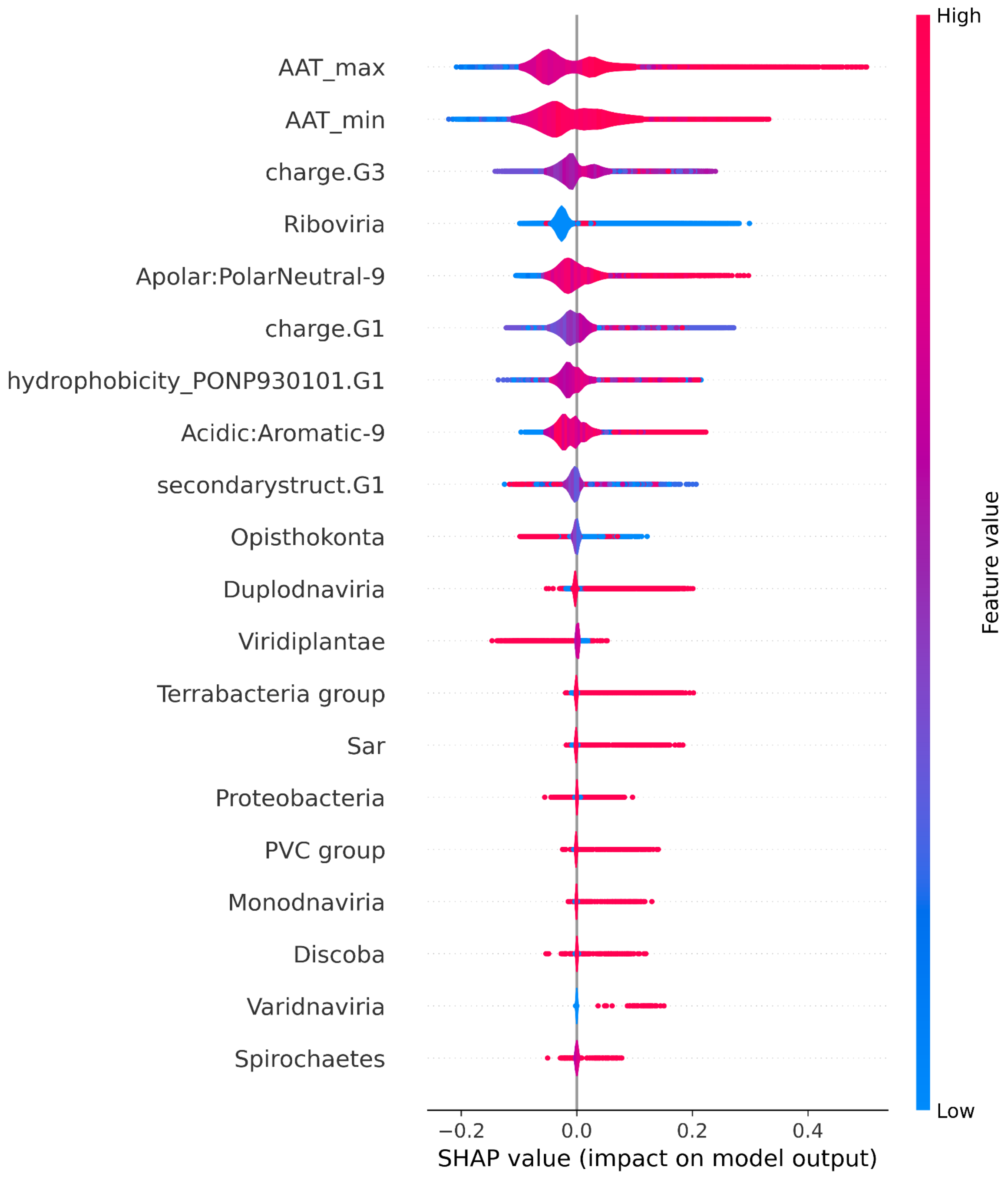


**Figure S2.** SHAP summary plot listing in descending order the 20 most important features to the Random Forest classifier considering the epitope1D training set, shown as coloured dots. The colouring scheme embodies the feature range values, from low in blue, to higher values in red colour. The X axis represents the SHAP values (values greater than 0 favour the epitope class and below this threshold contribute to the non-epitope class).

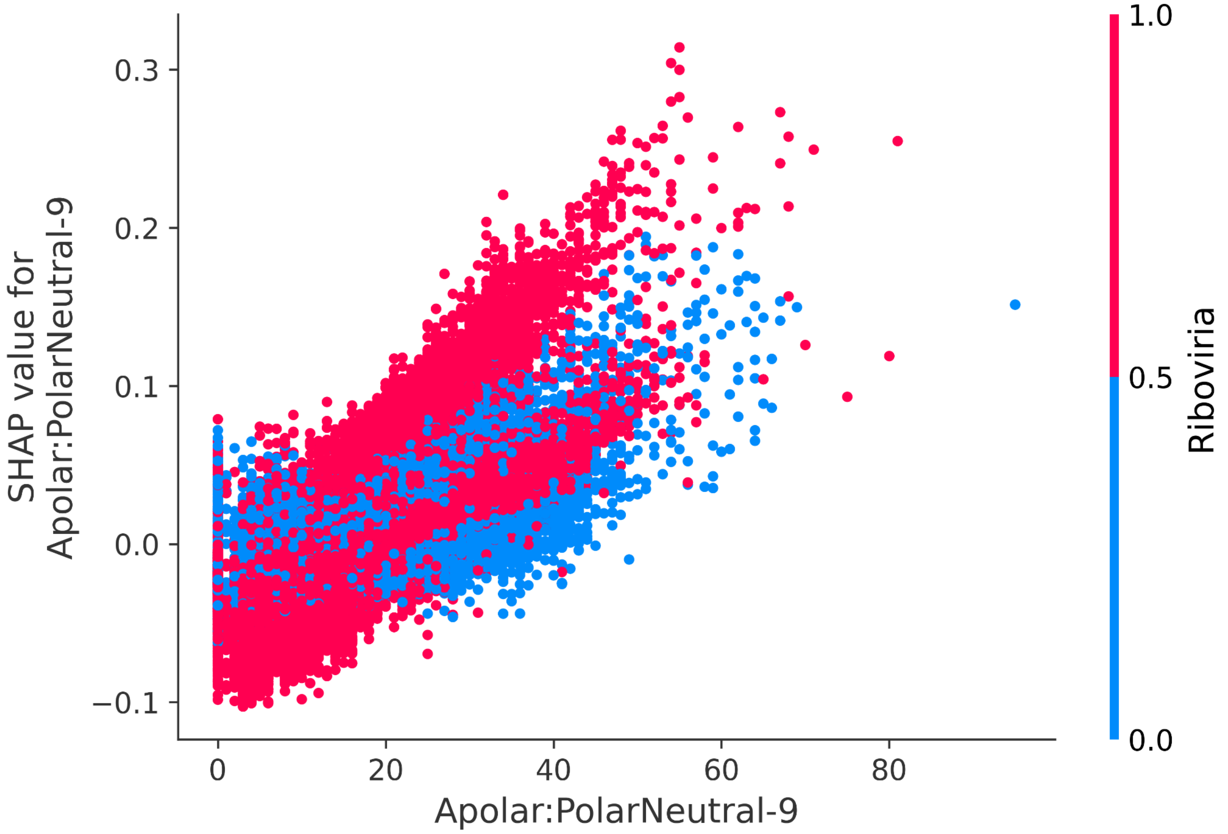


**Figure S3.** SHAP dependence plot showing the marginal effect of the feature Apolar:PolarNeutral-9 (values in the X axis) to its most correlated, the Riboviria Organism (symbolised in the colouring pattern). The Y axis represents the SHAP values itself, values above 0 contribute to the model decision into the epitope class and below to the non-epitope.

### TABLES

**Table S1.** Description of all applied features in the machine learning model. The Feature Class describes the basic name of the feature, while the Description briefly outlines its function. The Attributes column specifies the layout used for the features and the Value Type shows the data output type of each descriptor.

| Feature Class | Description | Attributes | Value Type |
|---|---|---|---|
| Amino Acid Composition using iFeatures. | Common compositions calculated solely on the peptide sequence. | k-spaced amino acid pairs;Dipeptides;Tripeptides;Enhanced and Grouped composition. | Float. |
| | Composition/Transition/Distribution (CTD) patterns of physicochemical and structural properties divided in 3 groups (G1,G2,G3), according to the amino acid type. | (1) Hydrophobicity, (2) Normalised van der Waals volume, (3) Polarity, (4) Secondary structure and (5) Charge and (6) Solvent accessibility. | Float. |
| Antigenicity Scale | Ratio value between epitope and non-epitope class taking amino acid groups. | (1) AAP(amino acid pairs) and (2) AAT(amino acid triplets). | Ratio: Float. (normalised between -1 to +1) |
| Graph-based Signatures | Model distance patterns among residue pairs (nodes) at different distance cutoffs and output as a cumulative sum. | Graph-labels: (1) prediction scales and (2) physicochemical properties. | Cumulative sum: Integer. |
| Organism Taxonomy | NCBI organismal taxonomy of the higher parent level in the lineage. | Twenty possible classes: Metamonada, Discoba, Sar, Viridiplantae, Opisthokonta, Terrabacteria group, Proteobacteria, PVC group, Spirochaetes, FCB group, Thermodesulfobacteria, Fusobacteria, Riboviria, Duplodnaviria, Monodnaviria, Varidnaviria, Ribozyviria, Anelloviridae, Naldaviricetes, Adnaviria. | Binary. |

**Table S2.** Performance comparison amidst proceedings methods using data derived from iBCE-EL method (training and testing sets) and ABCPred 1, all in a blind-test approach. Their performances were also extracted from EpitopeVec publication.

| **METHOD** | **MCC** | **ROC-AUC** | **F1** | **ACCURACY** |
|---|---|---|---|---|
| *Data set: iBCE-EL training* | | | | |
| BepiPred | 0.076 | 0.556 | **0.540** | 0.538 |
| BepiPred-2.0 | 0.045 | 0.51 | 0.410 | 0.509 |
| EpiDope | 0.064 | 0.582 | 0.370 | 0.505 |
| EpitopeVec | 0.085 | **0.555** | 0.520 | 0.538 |
| **epitope1D** | **0.131** | 0.545 | 0.281 | **0.562** |
| *Data set: iBCE-EL testing* | | | | |
| iBCE-EL | **0.454** | **0.786** | **0.730** | **0.734** |
| BepiPred | 0.104 | 0.568 | 0.550 | 0.558 |
| BepiPred-2.0 | 0.065 | 0.486 | 0.430 | 0.516 |
| EpiDope | 0.049 | 0.595 | 0.370 | 0.505 |
| EpitopeVec | 0.095 | 0.571 | 0.530 | 0.542 |
| **epitope1D** | 0.092 | 0.531 | 0.253 | 0.594 |
| *Data set: ABCPred 1* | | | | |
| ABCPred | 0.466 | 0.794 | -[a] | 0.730 |
| AAP | 0.518 | 0.782 | -[a] | 0.731 |
| LBTope | -[a] | -[a] | -[a] | 0.579 |
| iBCE-EL | 0.112 | 0.588 | 0.420 | 0.527 |
| BepiPred | 0.158 | 0.624 | 0.570 | 0.577 |
| BepiPred-2.0 | -0.040 | 0.399 | 0.350 | 0.493 |
| EpiDope | 0.059 | 0.599 | 0.350 | 0.506 |
| EpitopeVec | **0.714** | **0.929** | **0.860** | **0.856** |

| | | | | |
|---|---|---|---|---|
| **epitope1D** | 0.667 | 0.823 | 0.798 | 0.823 |

[a] Metric unavailable in original publication.

**Table S3**. List of the 28 descriptors applied in the final model, including the organismal taxonomy classes that can be grouped by Graph-based signatures (1-2), AAT Antigenicity ratio (3-4), Composition features (5-8) and Organism taxonomy (9-28).

| Features | | | | | |
|---|---|---|---|---|---|
| **1** | Graph-based Signature (Acidic:Aromatic-9) | **11** | Sar | **21** | Riboviria |
| **2** | Graph-based Signature (Apolar:PolarNeutral-9) | **12** | Viridiplantae | **22** | Duplodnaviria |
| **3** | AAT maximum | **13** | Opisthokonta | **23** | Monodnaviria |
| **4** | AAT minimum | **14** | Terrabacteria group | **24** | Varidnaviria |
| **5** | Composition (hydrophobicity_PONP930101.G1) | **15** | Proteobacteria | **25** | Ribozyviria |
| **6** | Composition (secondarystruct.G1) | **16** | PVC group | **26** | Anelloviridae |
| **7** | Composition (charge.G1) | **17** | Spirochaetes | **27** | Naldaviricetes |
| **8** | Composition (charge.G3) | **18** | FCB group | **28** | Adnaviria |
| **9** | Metamonada | **19** | Thermodesulfobacteria | | |
| **10** | Discoba | **20** | Fusobacteria | | |
